# Gut Microbiota Shifts with Lifetime Duration of Oral Contraception Use

**DOI:** 10.1101/2025.09.09.675238

**Authors:** Gauri Paul, Sayumi York, Rosa Alcazar

## Abstract

To investigate the effects of lifetime duration of oral contraceptive (OC) use, we reanalyzed publicly available 16S rRNA sequencing data from fecal samples of OC users and non-users collected at different timepoints in the menstrual cycle. We profiled the microbial composition of subjects and identified 32 significantly differentially abundant Amplicon Sequence Variants (padj<0.05) with increasing lifetime use duration including 16 ASVs belonging to class Clostidia that decreased with longer OC use. Some Clostridia species are known to recycle estrogen; lifetime duration of OC exposure could thus influence women’s health through gut microbiome composition and hormone regulation.

## Description

Microbial communities are now recognized as beneficial and essential to the holobiont. Composition of the gut microbiome is influenced by a variety of factors including diet, stress, environment, age, gender, ethnicity, and lifetime medication exposure (Cresci & Bawden, 2015). The gut microbiota has been of particular interest to the scientific community for its role in immunity, metabolism, disease onset, and behavior (Young, 2017). In addition, the gut microbiome can interact with sex hormones, influencing and being influenced by estrogen metabolism (He et al,. 2021), which makes it relevant to studies of women’s health.

Historically, there has been a disparity in the number of studies conducted on female subjects, especially on issues that are gender-specific or common in women such as osteoporosis, anxiety, and heart disease (Siddiqui et al., 2022). Dysbiosis of the gut and vaginal microbiome has also been linked to conditions such as polycystic ovarian syndrome (PCOS), cancer, menopause, and post-menopausal ailments (Siddiqui et al., 2022). Previous studies indicate hormonal contraceptive use, combined with the progression of the menstrual cycle, can alter the female microbiome across different body sites (Krog et al., 2022). Researchers have confirmed that the hormonal environment can influence the gut microbiome. Conversely, the gut microbiome can regulate sex hormones, particularly estrogen, through metabolites, immune modulation and inflammation (He et al., 2021).

Hormonal oral contraceptives (OC) are used by women of reproductive age, both to prevent pregnancy and treat conditions such as hormonal acne, PCOS, endometriosis, and menstrual pain (National Research Council (US) Committee on Population, 1989). Studies have shown that species richness and diversity of the gut microbiome have a bidirectional relationship with systemic estrogen levels (Flores et al., 2012). It is suggested that use of estrogen-progestin based OC has been linked to an elevation in breast cancer risk (Kim & Munster, 2025). These connections highlight the need to study how OC use may alter microbiome composition and estrogen regulation.

In a recent study published by Terrazas, et al. overall microbial diversity did not differ between OC users and non-users on day 2 of the menstrual cycle, but significant cycle-dependent effect of OCs on species richness was observed on day 21 (Terrazas et al., 2025). Although data on subjects’ lifetime duration of OC use was collected, this study did not present any analysis relating to the length of time on OC to the gut microbiota. Additionally, the composition of the gut microbiome at the level of individual samples was not reported. Here, we reanalyzed Terrazas et al.’s dataset to profile gut microbiota composition at individual samples and test whether lifetime duration of OC use was associated with differential abundance of taxa.

Terrazas et al.’s original study analyzed 16S rRNA sequencing data collected from the fecal samples of OC users and non-users at two timepoints (day 2 and day 21) in the menstrual cycle. Non-users were defined as having no OC use for a minimum of 3 months prior to the start of the study; however, this does not disqualify prior use of OC. Due to a lack of follow-up in some subjects, there were more samples collected for both users and non-users on day 2 (n=45 and n=19 respectively) compared to day 21 (n=39 and n=17 respectively). A total of 5457 Amplicon Sequence Variants (ASVs) were recovered across all samples.

We found that the most abundant bacterial phylum in the majority of subjects were Bacteroidota, Firmicutes, and Proteobacteria (Figure 1A). Only Bacteroidota and Firmicutes were detected in all samples. Phyla Actinobacteriota and Proteobacteria were present in nearly all (98.3%) samples, Desulfobacterota in 77.4%, Verrucomicrobiota in 48.7% of samples. The mean relative abundance of phylum Bacteroidota across all samples was 52.91±14.74%, ranging from 1.89% to 85.35%. Phylum Firmicutes had a mean relative abundance of 41.06±13.66%, ranging from 11.74% to 97.70%. The mean relative abundance of phylum Proteobacteria was 3.43±4.53%, ranging from 0.00% to 27.99%.

**Figure 1.**
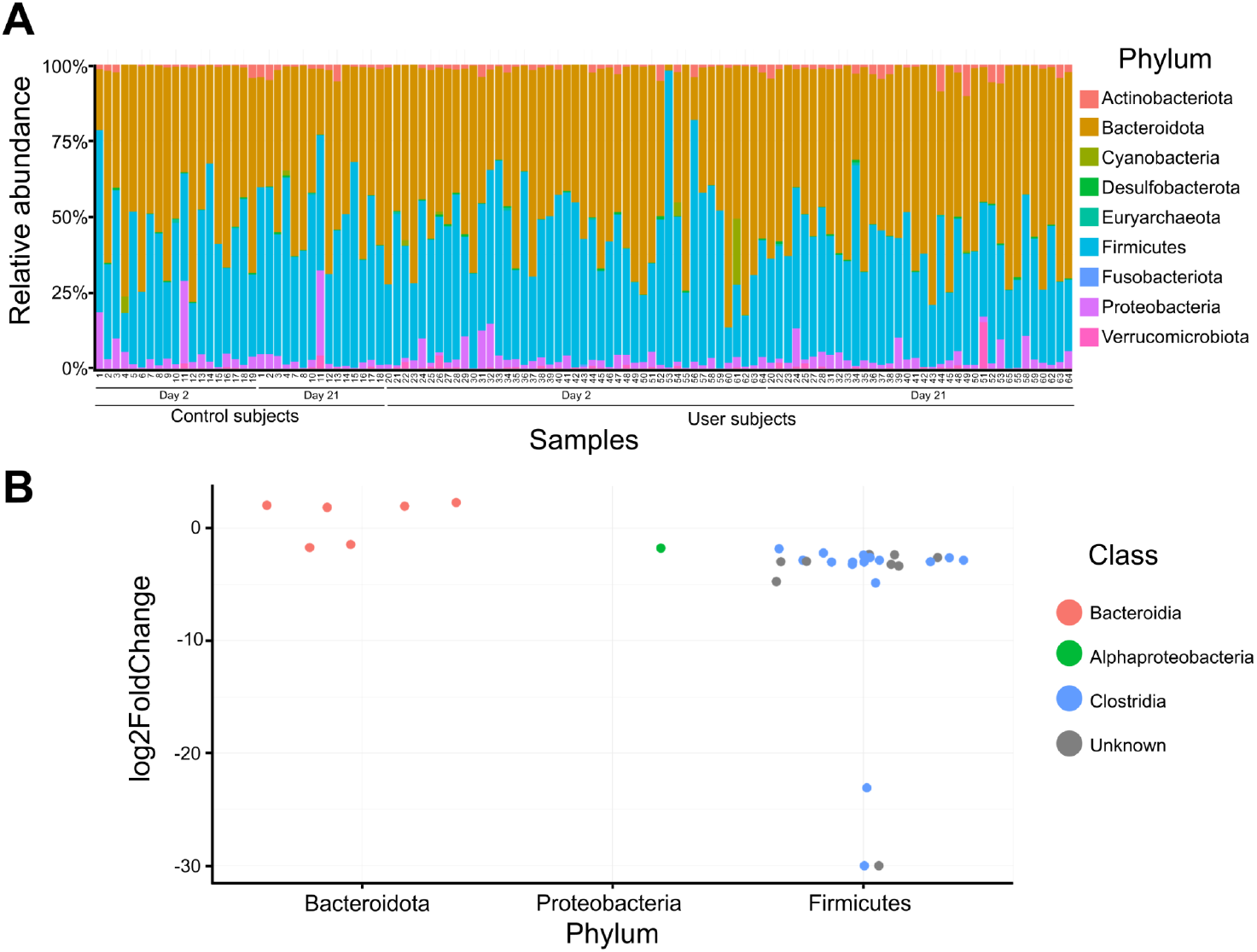
**A. Relative Abundance plot showing the gut microbial composition for oral contraceptive (OC) users and non-users at different time points in the menstrual cycle.** It represents major phyla like Firmicutes, Bacteroidota and Bacteroidota composing the human gut microbiota in all subjects, and comparing OC users and controls on Day 2 and Day 21 of the menstrual cycle. **B. Differential Abundance of ASVs across all samples with increasing lifetime OC use**. The plot shows log2FoldChange (L2FC) values for significantly differentially abundant Amplicon Sequence Variants (ASVs). Positive values indicate taxa that increase in abundance with each year of lifetime OC use. Most significant decreases were found in class Clostridia belonging to phylum Firmicutes.

Three atypical subjects were also identified whose microbial composition did not follow the general pattern of relative abundance. Subject 11 (non-user) showed a reduced abundance of Bacteriodota and the highest relative abundance of Proteobacteria compared to all other subjects. This subject had a Bacteroidota relative abundance of 34.71%, Firmicutes of 35.48%, and Proteobacteria of 27.15% on day 2 and 21.65.%, 44.60%, and 27.99% respectively, on day 21 of the menstrual cycle. Subject 53 (user) showed the highest relative abundance of Firmicutes across all samples: 97.69% on day 2, reduced to 31.00% on day 21. Lastly, subject 61 (user), was the only sample with a high abundance of Cyanobacteria (21.88%).

To compare the effects of total years a subject has been on OC use across their lifetime, differential abundance analysis was performed on a subset of samples with available data. In this analysis, 8 non-users had prior OC use, with an average lifelong duration of 2.77 ± 2.3 years of OC use while 43 users had an average lifelong duration of 3.28 ± 2.31 years. 32 significantly differentially abundant (padj<0.05) ASVs were identified in three phyla: 6 Bacteroidota, 25 Firmicutes, and 1 Proteobacteria ASVs (Figure 1B). Out of the 32 ASVs, 4 had a positive Log2FoldChange (L2FC) and 28 had negative L2FC values. The 4 ASVs with a positive L2FC within the phylum Bacteroidota were identified as *Bacteroides plebeius;* the remaining 2 ASVs belonging to genus *Provotella* had negative L2FC. All 25 ASVs belonging to phylum Firmicutes had negative L2FC values. 16 out of 25 Firmicutes ASVs belonged to class Clostridia.

Firmicutes, Bacteroidota, and Proteobacteria were the major components of gut communities and ASVs from these groups significantly changed with increasing duration of lifetime OC use. Previous studies have found that Bacteroidota and Firmicutes are two of the most abundant phyla in the human gut microbiota (Magne et al., 2020; Rinninella et al., 2019). The Firmicutes/Bacteroidota ratio has been used as a marker of dysbiosis in a large number of microbiome studies including those relating to obesity, type 2 diabetes, cardiovascular conditions, and cancers, with an increasing Firmicutes/Bacteroidota ratio and reduced bacterial diversity linked to dysbiosis and inflammation, thought to be the underlying cause of many of these conditions (Baker et al., 2017).

Our study found 16 ASVs belonging to class Clostridia that significantly decreased in abundance with increasing lifetime OC use duration. While our methods were unable to annotate to the level of genus, our results support Terrazas et al.’s original finding that *Clostridium* abundance was lower in OC users compared to non-users. Some species of *Clostridium* produce beta-glucuronidase, an enzyme that enables enterohepatic recirculation of estrogen, regulating its bioavailability and contributing to hormonal homeostasis (Escorcia Mora et al., 2025). Reduction in these bacteria in the gut may disrupt reproductive function, while overrepresentation of Clostridium in the gut has been linked to conditions such as endometriosis, breast cancer, and infertility (Escorcia Mora et al., 2025).

The reversion to a baseline menstrual cycle can take up to 9 months following termination of OC use (Gnoth et al., 2002). In Terrazas et al.’s study, the non-users included subjects who refrained from OCs for a minimum of 3 months. As a result, the control group may have included subjects who had used OCs within the previous 9 months and whose cycles, hormone levels, and gut microbiome were still returning to a baseline state. Future research should explore the longevity of the effect of OC on the gut microbiome and clarify how long these microbial effects persist after discontinuation.

Our study focused on monophasic estrogen pills, but other oral contraceptive types, such as combination birth control pills and progestin-only pills, are also widely used (Hall & Trussell, 2012). Other methods of contraception, including Long-Acting Reversible Contraceptives (LARCS) such as copper and hormone-based intrauterine devices (IUDs) and implants are gaining popularity, especially in adolescents and can also be analyzed for impact on the gut microbiome and women’s health in future (Durante et al., 2023). Millions of women take oral contraceptives regularly (Haakenstad et al., 2022). Understanding the long-term effects of its prolonged OC use on the gut microbiome is therefore an important health priority. Ideally, future work should also incorporate the effects of discontinuous OC use and directly compare the microbiome of the same individual before and during OC use to control for other contributing factors to microbial composition such as intra-individual variation (Guthrie et al., 2022).

In summary, we reanalyzed Terrazas et al.’s 2025 dataset to examine whether lifetime duration of OC use influences the gut microbiome. We confirmed that Firmicutes, Bacteroidota, and Proteobacteria dominate the microbial community, and also identified 32 differentially abundant ASVs, including 16 Clostridia that decreased with longer OC use. These findings highlight the importance of considering duration of exposure, not just the presence or absence of use, when studying the effects of oral contraceptives. They also suggest that long-term OC use could gradually reshape the gut microbiome. This raises new questions about how such changes could influence estrogen metabolism and hormone regulation.

## Methods

Data from Terrazas et al. (2025) were downloaded from the authors’ Github (https://github.com/fernanda-t/microbiome-oc). Microbial composition was reanalyzed and visualized using the R packages phyloseq and ggplot2 (Ihaka & Gentleman, 1996; McMurdie & Holmes, 2013; Terrazas et al., 2025; Wickham, 2011). To examine the effect of the duration of continuous OC use on the gut microbiota we examined a subset of samples for which this data was available which contained both (n=16) non-users and (n=73) OC users. Differential abundance of ASVs related to the number of years on oral contraceptives regardless of dosage was assessed using DESeq2 using the alternative geometric mean to estimate size factors with a *p*-adj cutoff of 0.05 (Love et al., 2014).

## Acknowledgements

This work was supported by NIH grant UE5HG013799-01. We thank the C-MOOR team for technical assistance and to fellow C-MOOR Scholars Gugulethu Sakana, Grace Ekalle, and Madeleine Gerard for their continued support throughout this project.

## CRediT

**Gauri Paul:** Conceptualization, formal analysis, investigation process, software, visualization, writing (original draft)

**Sayumi York:** Conceptualization, formal analysis, investigation, methodology, project administration, resources, software, supervision, visualization, writing (original draft)

**Rosa Alcazar**: Supervision, writing (review and editing)

